# Systematic interrogation of diverse Omic data reveals interpretable, robust, and generalizable transcriptomic features of clinically successful therapeutic targets

**DOI:** 10.1101/220848

**Authors:** Andrew D. Rouillard, Mark R. Hurle, Pankaj Agarwal

## Abstract

Target selection is the first and pivotal step in drug discovery. An incorrect choice may not manifest itself for many years after hundreds of millions of research dollars have been spent. We collected a set of 332 targets that succeeded or failed in phase III clinical trials, and explored whether Omic features describing the target genes could predict clinical success. We obtained features from the recently published comprehensive resource: Harmonizome. Nineteen features appeared to be significantly correlated with phase III clinical trial outcomes, but only 4 passed validation schemes that used bootstrapping or modified permutation tests to assess feature robustness and generalizability while accounting for target class selection bias. We also used classifiers to perform multivariate feature selection and found that classifiers with a single feature performed as well in cross-validation as classifiers with more features (AUROC=0.57 and AUPR=0.81). The two predominantly selected features were mean mRNA expression across tissues and standard deviation of expression across tissues, where successful targets tended to have lower mean expression and higher expression variance than failed targets. This finding supports the conventional wisdom that it is favorable for a target to be present in the tissue(s) affected by a disease and absent from other tissues. Overall, our results suggest that it is feasible to construct a model integrating interpretable target features to inform target selection. We anticipate deeper insights and better models in the future, as researchers can reuse the data we have provided to improve methods for handling sample biases and learn more informative features. Code, documentation, and data for this study have been deposited on GitHub at https://github.com/arouillard/omic-features-successful-targets.

**AUTHOR SUMMARY:** Drug discovery often begins with a hypothesis that changing the abundance or activity of a target—a biological molecule, usually a protein—will cure a disease or ameliorate its symptoms. Whether a target hypothesis translates into a successful therapy depends in part on the characteristics of the target, but it is not completely understood which target characteristics are important for success. We sought to answer this question with a supervised machine learning approach. We obtained outcomes of target hypotheses tested in clinical trials, scoring targets as successful or failed, and then obtained thousands of features (i.e. properties or characteristics) of targets from dozens of biological datasets. We statistically tested which features differed between successful and failed targets, and built a computational model that used these features to predict success or failure of targets in clinical trials. We found that successful targets tended to have more variable mRNA abundance from tissue to tissue and lower average abundance across tissues than failed targets. Thus, it is probably favorable for a target to be present in the tissue(s) affected by a disease and absent from other tissues. Our work demonstrates the feasibility of predicting clinical trial outcomes from target features.

## INTRODUCTION

More than half of drug candidates that advance beyond phase I clinical trials fail due to lack of efficacy (1, 2). One possible explanation for these failures is sub-optimal target selection (3). Many factors must be considered when selecting a target for drug discovery (4, 5). Intrinsic factors include the likelihood of the target to be tractable (can the target’s activity be altered by a compound, antibody, or other drug modality?), safe (will altering the target’s activity cause serious adverse events?), and efficacious (will altering the target’s activity provide significant benefit to patients?). Extrinsic factors include the availability of investigational reagents and disease models for preclinical target validation, whether biomarkers are known for measuring target engagement or therapeutic effect, the duration and complexity of clinical trials required to prove safety and efficacy, and the unmet need of patients with diseases that might be treated by modulating the target.

Over the past decade, technologies have matured enabling high-throughput genome-, transcriptome-, and proteome-wide profiling of cells and tissues in normal, disease, and experimentally perturbed states. In parallel, researchers have made substantial progress curating or text-mining biomedical literature to extract and organize information about genes and proteins, such as molecular functions and signaling pathways, into structured datasets. Taken together, both efforts have given rise to a vast amount of primary, curated, and text-mined data about genes and proteins, which are stored in online repositories and amenable to computational analysis (6, 7).

To improve the success rate of drug discovery projects, researchers have investigated whether any features of genes or proteins are useful for target selection. These computational studies can be categorized according to whether the researchers were trying to predict tractability (8, 9), safety (10-13), efficacy (no publications to our knowledge), or overall success (alternatively termed “drug target likeness”) (8, 13-26). Closely related efforts include disease gene prediction, where the goal is to predict genes mechanistically involved in a given disease (27-32), and disease target prediction, where the goal is to predict genes that would make successful drug targets for a given disease (33-35).

To our knowledge, we report the first screen for features of genes or proteins that distinguish targets of approved drugs from targets of drug candidates that failed in clinical trials. In contrast, related prior studies have searched for features that distinguish targets of approved drugs from the rest of the genome (or a representative subset) (13, 15-25). Using the remainder of the genome for comparison has been useful for finding features enriched among successful targets, but it is uncertain whether these features are specific to successful targets or are enriched among targets of failed drug candidates as well. Our study aims to fill this knowledge gap by directly testing for features that separate targets by clinical outcome, expanding the scope of prior studies that have investigated how genetic disease associations (36) and publication trends (37) of targets correlate with clinical outcome.

Our work has five additional innovative characteristics. First, we included only targets of drugs that are presumed to be selective (no documented polypharmacology) to reduce ambiguity in assigning clinical trial outcomes to targets. Second, we included only phase III failures to enrich for target efficacy failures, as opposed to safety and target engagement failures, which are more common in phase I and phase II (2). Third, we excluded targets of assets only indicated for cancer, as studies have observed that features of successful targets for cancer differ from features of successful targets for other indications (22, 23), moreover, cancer trials fail more frequently than trials for other indications (2). Fourth, we interrogated a diverse and comprehensive set of features, over 150,000 features from 67 datasets covering 16 feature types, whereas prior studies have examined only features derived from protein sequence (16-18, 24, 25), protein-protein interactions (13, 15, 18-23), Gene Ontology terms (13, 15, 16), and gene expression profiles (15, 19, 21, 25). Fifth, because targets of drugs and drug candidates do not constitute a random sample of the genome, we implemented a suite of tests to assess the robustness and generalizability of features identified as significantly separating successes from failures in the biased sample.

A handful of the initial 150,000+ features passed our tests for robustness and generalizability to new targets or target classes. Interestingly, these features were predominantly derived from gene expression datasets. *Notably, two significant features were discovered repeatedly in multiple datasets: successful targets tended to have lower mean mRNA expression across tissues and higher expression variance than failed targets.* We also trained a classifier to predict phase III success probabilities for untested targets (no phase III clinical trial outcomes reported for drug candidates that selectively modulate these targets). We identified 943 targets with sufficiently unfavorable expression characteristics to be predicted twice as likely to fail in phase III clinical trials as past phase III targets. Furthermore, we identified 2,700,856 target pairs predicted with 99% consistency to have a 2-fold difference in success probability. Such pairwise comparisons may be useful for prioritizing short lists of targets under consideration for a therapeutic program. We conclude this paper with a discussion of the biases and limitations faced when attempting to analyze, model, or interpret data on clinical trial outcomes.

## RESULTS

### Examples of successful and failed targets obtained from phase III clinical trial reports

We extracted phase III clinical trial outcomes reported in Pharmaprojects (38) for drug candidates reported to be selective (single documented target) and tested as treatments for non-cancer diseases. We grouped the outcomes by target, scored targets with at least one approved drug as successful (N_S_=259), and scored targets with no approved drugs and at least one documented phase III failure as failed (N_F_=72) (Supplementary Table S1). The target success rate (77%) appears to be inflated relative to typically reported phase III success rates (58%) (2) because we scored targets by their best outcome across multiple trials.

### Comprehensive and diverse collection of target features obtained from the Harmonizome

We obtained target features from the Harmonizome (39), a recently published collection of features of genes and proteins extracted from over 100 Omics datasets. We limited our analysis to 67 datasets that are in the public domain or GSK had independently licensed (Table 1). Each dataset in the Harmonizome is organized into a matrix with genes labeling the rows and features such as diseases, phenotypes, tissues, and pathways labeling the columns. We included the mean and standard deviation calculated along the rows of each dataset as additional target features. These summary statistics provide potentially useful and interpretable information about targets, such as how many pathway associations a target has or how variable a target’s expression is across tissues.

**Table 1.**
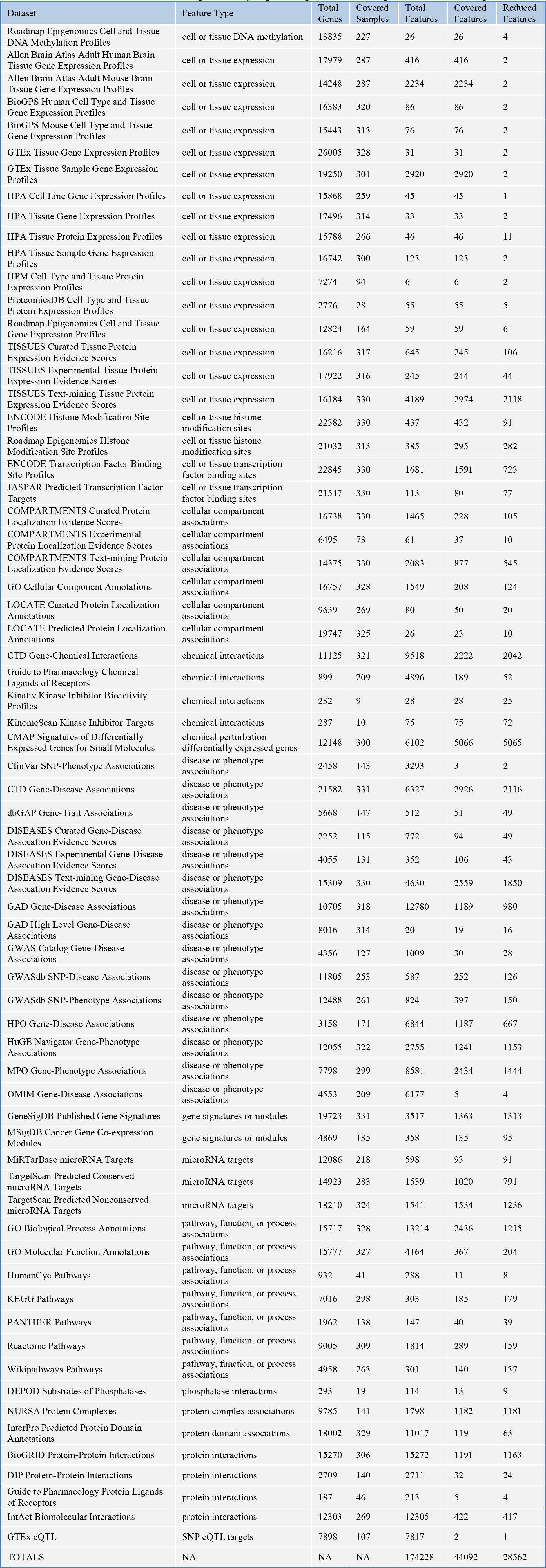
Datasets tested for features significantly separating successful targets from failed targets.

The datasets contained a total of 174,228 features covering 16 feature types (Table 1). We restricted our analysis to 44,092 features that had at least three non-zero values for targets assigned a phase III outcome. Many datasets had strong correlations among their features. To reduce feature redundancy and avoid excessive multiple hypothesis testing while maintaining interpretability of features, we replaced each group of highly correlated features with the group mean feature and assigned it a representative label (Fig 1, Supplementary Table S2). The number of features shrunk to 28,562 after reducing redundancy.

**Fig 1.**
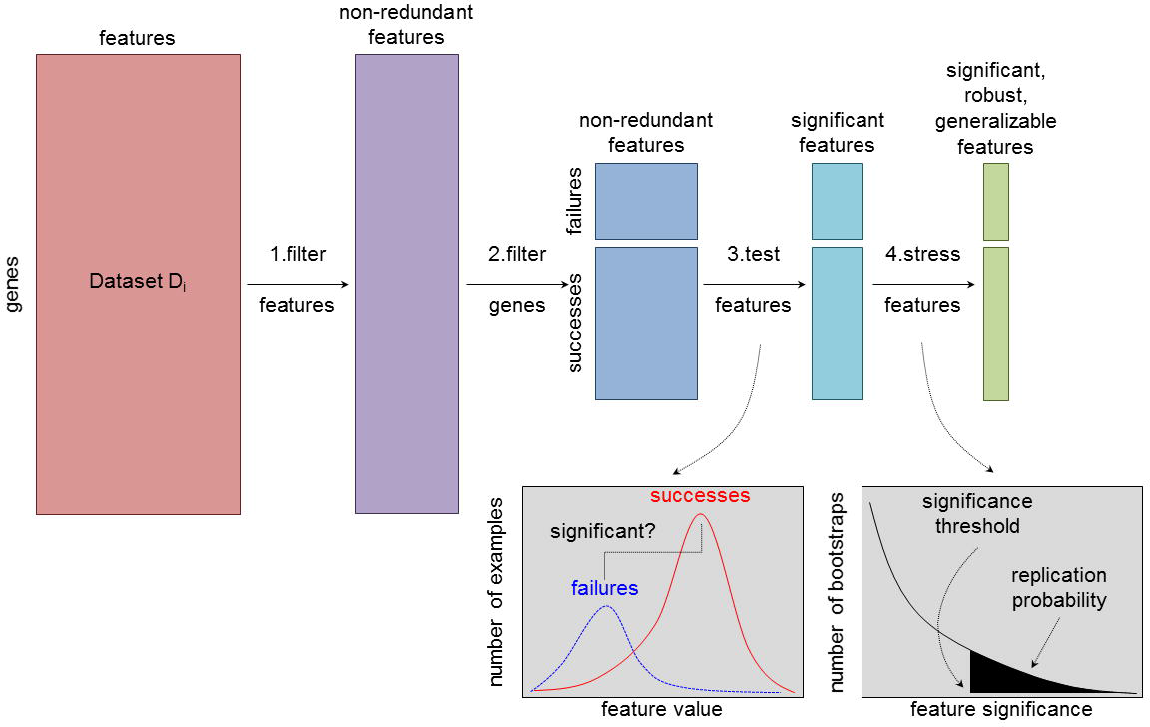
Feature Selection Pipeline. Each dataset took the form of a matrix with genes labeling the rows and features labeling the columns. We appended the mean and standard deviation computed across all features as two additional features. **Step 1:** We filtered the columns to eliminate redundant features, replacing each group of correlated features with the group average feature, where a group was defined as features with squared pair-wise correlation coefficient r^2^ ≥ 0.5. If the dataset mean feature was included in a group of correlated features, we replaced the group with the dataset mean. **Step 2:** We filtered the rows for targets with clinical trial outcomes of interest: targets of selective drugs approved for non-cancer indications (successes) and targets of selective drug candidates that failed in phase III clinical trials for non-cancer indications (failures). **Step 3:** We tested the significance of each feature as an indicator of success or failure using permutation tests to quantify the significance of the difference between the means of the successful and failed targets. We corrected for multiple hypothesis testing using the Benjamini-Yekutieli method to control the false discovery rate at 0.05 within each dataset. **Step 4:** We “stressed” the significant features with additional tests to assess their robustness and generalizability. For example, we used bootstrapping to estimate probabilities that the significance findings will replicate on similar sets of targets.

### Target features tested for correlation with phase III outcome

We performed permutation tests (40, 41) on the remaining 28,562 target features to find features with a significant difference between the successful and failed targets, and we corrected p-values for multiple hypothesis testing using the Benjamini-Yekutieli method (42) (Fig 1, Supplementary Table S2). We used permutation testing to apply the same significance testing method to all features, since they had heterogeneous data distributions. We detected 19 features correlated with clinical outcome at a within-dataset false discovery rate of 0.05 (Table 2). The significant features were derived from 7 datasets, of which 6 datasets were gene expression atlases: Allen Brain Atlas adult human brain tissues (43, 44), Allen Brain Atlas adult mouse brain tissues (43, 45), BioGPS human cell types and tissues (46-48), BioGPS mouse cell types and tissues (46-48), Genotype-Tissue Expression Project (GTEx) human tissues (49, 50), and Human Protein Atlas (HPA) human tissues (51). The remaining dataset, TISSUES (52), was an integration of experimental gene and protein tissue expression evidence from multiple sources. Two correlations were significant in multiple datasets: successful targets tended to have lower mean expression across tissues and higher expression variance than failed targets.

**Table 2.**
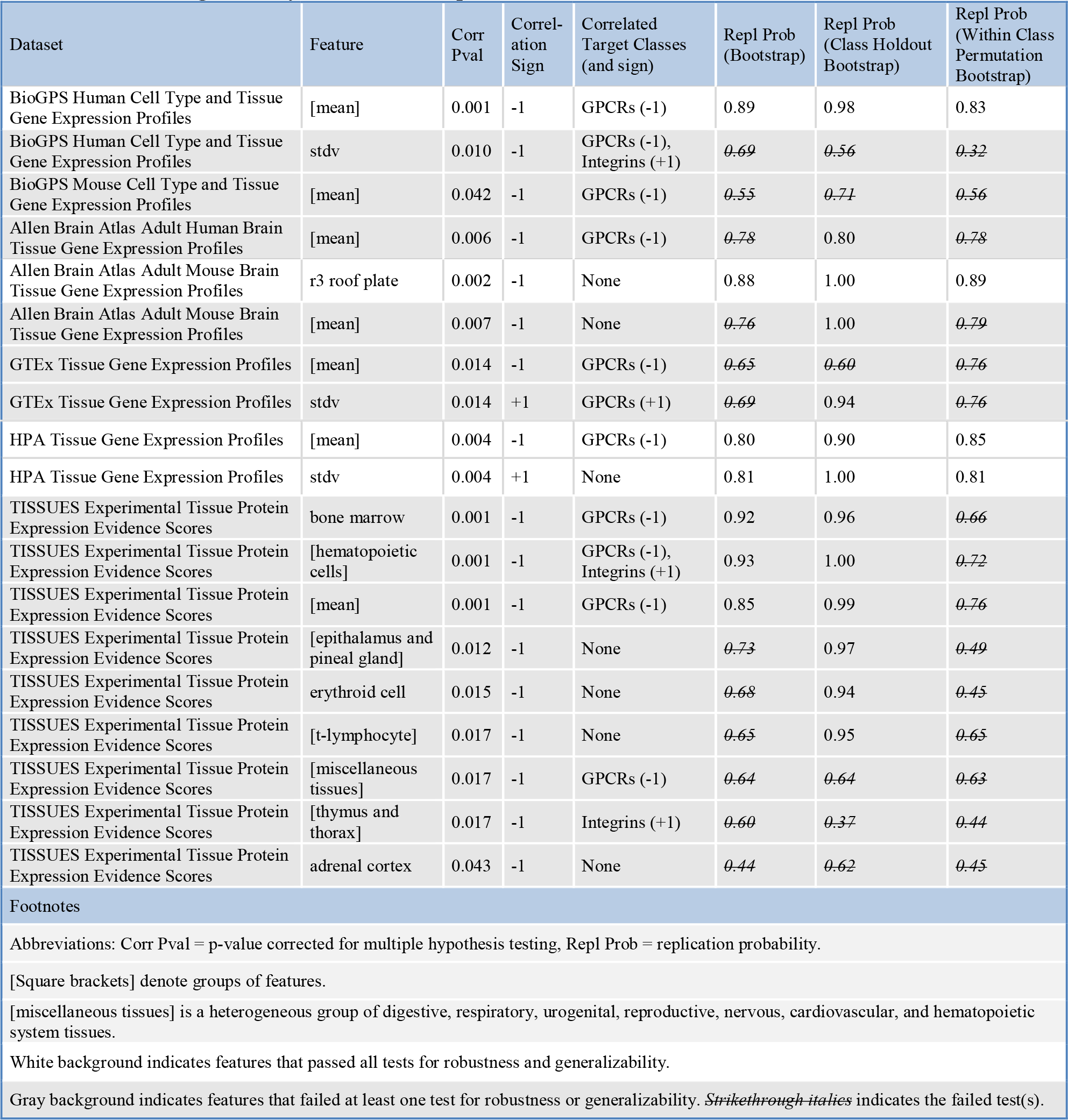
Features significantly correlated with phase III outcome.

### Significant features tested for robustness to sample variation and generalization across target classes

Because targets of drugs and drug candidates do not constitute a random sample of the genome, features that separate successful targets from failed targets in our sample may perform poorly as genome-wide predictors of success versus failure. We performed three analyses to address this issue (Fig 1).

#### Robustness to sample variation

We used bootstrapping (53, 54) (sampling with replacement from the original set of examples to construct sets of examples equal in size to the original set) to investigate how robust our significance findings were to variation in the success and failure examples. For each dataset that yielded significant features in our primary analysis, we repeated the analysis on 1000 bootstrap samples and quantified the replication probability (55) of each feature as the fraction of bootstraps yielding a significant correlation with phase III outcome at a within-dataset false discovery rate of 0.05. Twelve features had less than 80% probability (considered a strong replication probability in (55)) that their correlation with clinical outcome will generalize to new examples (Table 2).

#### Robustness to target class variation

We tested if any of the significance findings depended upon the presence of targets from a single target class in our sample. We obtained target class labels (i.e. gene family labels) from the HUGO Gene Nomenclature Committee (56), tested if any target classes were significantly correlated with phase III outcome, and then tested if these classes were correlated with any features. The GPCR and integrin classes were correlated with phase III outcome as well as several features (Table 2). This raised the possibility that instead of these features being genome-wide indicators of clinical outcome, they were simply reflecting the fact that many GPCRs have succeeded (62/70, p<0.05) or that integrins have failed (3/3, p<0.01). To test this possibility, we repeated the bootstrapping procedure described above to obtain replication probabilities, except excluded GPCRs and integrins from being drawn in the bootstrap samples. Six features had less than 80% probability that their correlation with clinical outcome will generalize to new target classes (Table 2).

#### Generalization across target classes

In the preceding analysis, we checked one target class at a time for its impact on our significance findings. To broadly test whether features generalize across target classes, we repeated the permutation testing described in our initial analysis, but only shuffled the success/failure labels within target classes, inspired by the work of Epstein et al. (57) on correcting for confounders in permutation testing. By generating a null distribution with preserved ratio of successes to failures within each target class, features must correlate with clinical outcome within multiple classes to be significant, while features that discriminate between classes will not be significant. We repeated the modified permutation tests on 1000 bootstrap samples to obtain replication probabilities. We rejected fifteen features that had less than 80% probability that their correlation with clinical outcome generalizes across target classes (Table 2). This set of fifteen features included all features with less than 80% replication probability in either of the previous two tests. The remaining robust and generalizable features were: 1) mean mRNA expression across tissues (HPA and BioGPS human tissue expression datasets), 2) standard deviation of expression across tissues (HPA human tissue expression dataset), and 3) expression in r3 roof plate (Allen Brain Atlas adult mouse brain tissue expression dataset). The r3 roof plate expression profile was correlated with mean expression across tissues in the Allen Brain Atlas dataset (r^2^=0.47), falling just below the r^2^=0.5 cut-off that would have grouped r3 roof plate with the mean expression profile during dimensionality reduction.

### Classifier-based assessment of feature usefulness and interpretability

Statistical significance did not guarantee the remaining features would be useful in practice for discriminating between successes and failures. To test their utility, we trained a classifier to predict target success or failure, using cross-validation to select a model type (Random Forest or logistic regression) and a subset of features useful for prediction. Because we used all targets with phase III outcomes for the feature selection procedure described above, simply using the final set of features to train a classifier on the same data would yield overly optimistic performance, even with cross-validation. Therefore, we implemented a nested cross-validation routine to perform both feature selection and model selection (58).

#### Cross-validation routine

The outer loop of the cross-validation routine had five steps (Fig 2): 1) separation of targets with phase III outcomes into training and testing sets, 2) univariate feature selection using the training set, 3) aggregation of features from different datasets into a single feature matrix, 4) classifier-based feature selection and model selection using the training set, and 5) evaluation of the classifier on the test set. Step 4 used an inner loop with 5-fold cross-validation repeated 20 times to estimate the performance of different classifier types (Random Forest or logistic regression) and feature subsets (created by incremental feature elimination). The simplest classifier (least number of features, with logistic regression considered simpler than Random Forest) with cross-validation values for area under the receiver operating characteristic curve (AUROC) and area under the precision-recall curve (AUPR) within 95% of maximum was selected. The outer loop used 5-fold cross-validation repeated 200 times, which provided 1000 train-test cycles for estimating the generalization performance of the classifier and characterizing the consistency of the selected features and model type.

**Fig 2.**
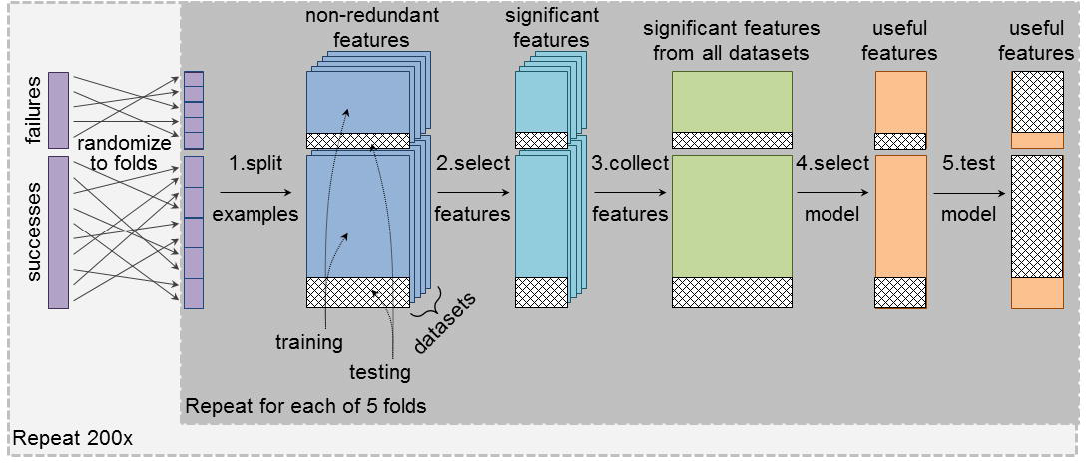
Modeling Pipeline. We trained a classifier to predict phase III clinical trial outcomes, using 5-fold cross-validation repeated 200 times to assess the stability of the classifier and estimate its generalization performance. For each fold of cross-validation, modeling began with the non-redundant features for each dataset. **Step 1:** We split the targets with phase III outcomes into training and testing sets. **Step 2:** We performed univariate feature selection using permutation tests to quantify the significance of the difference between the means of the successful and failed targets in the training examples. We controlled for target class as a confounding factor by only shuffling outcomes within target classes. We accepted features with adjusted p-values less than 0.05 after correcting for multiple hypothesis testing using the Benjamini-Yekutieli method. **Step 3:** We aggregated significant features from all datasets into a single feature matrix. **Step 4:** We performed incremental feature elimination with an inner 5-fold cross-validation loop repeated 20 times to select the type of classifier (Random Forest or logistic regression) and smallest subset of features that had cross-validation area under the receiver operating characteristic curve (AUROC) and area under the precision-recall curve (AUPR) values within 95% of maximum. **Step 5:** We refit the selected model using all the training examples and evaluated its performance on the test examples.

#### Classifier consistency

Simple models were consistently selected for the classifier (Table 3, Supplementary Table S3). In 1000 train-test cycles, a logistic regression model with one feature was selected most the time (66%), followed in frequency by a logistic regression model with two features (8%), a Random Forest model with two features (8%), and a logistic regression model with three features (6%). Other combinations of model type (logistic regression or Random Forest) and number of features (ranging from 1 to 8) appeared 11% of the time (each 4% or less). For one of the train-test cycles (0.1%), no significant features were found in the univariate feature selection step, resulting in a null model. Note that the logistic regression models were selected primarily because we imposed a preference for simple and interpretable models, not because they performed better than Random Forest models. The Random Forest model tended to perform as well as the logistic regression model on the inner cross-validation loop, with AUROC = 0.62 ± 0.06 for Random Forest and 0.63 ± 0.05 for logistic regression (Supplementary Table S4).

**Table 3.**
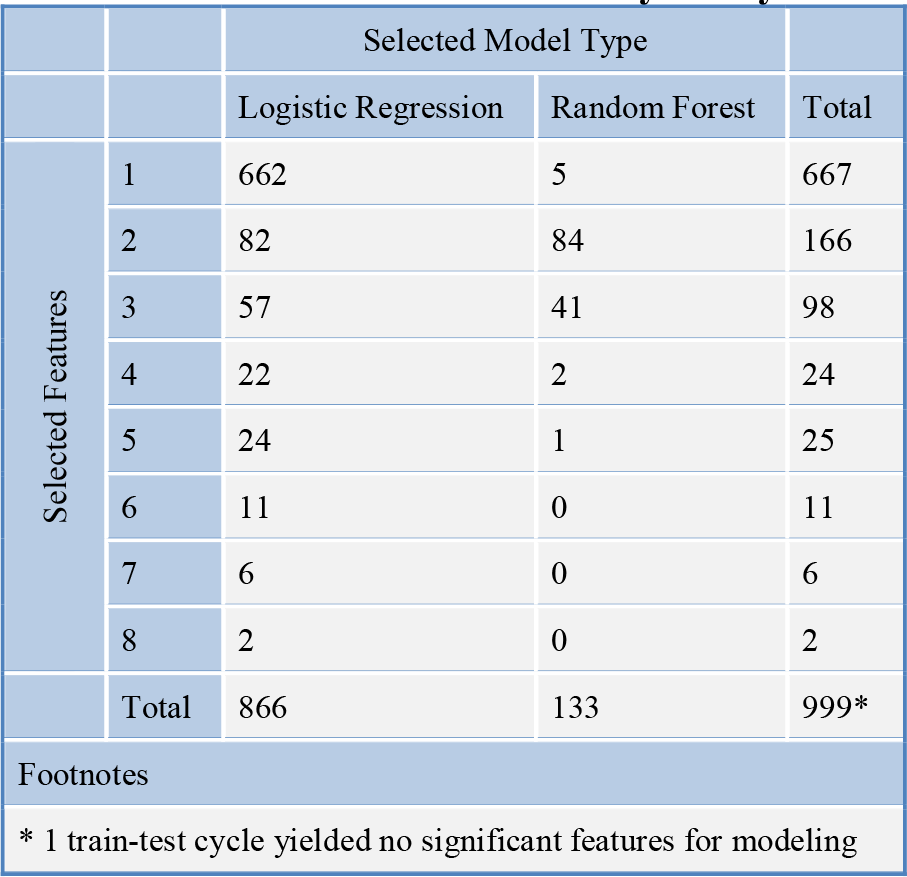
Distribution of train-test cycles by classifier type and number of selected features.

Gene expression features were consistently selected for the classifier (Table 4, Supplementary Table S3). Mean mRNA expression across tissues and standard deviation of expression across tissues had frequencies of 69% and 59%, respectively. More precisely, 36% of the models used mean mRNA expression across tissues as the only feature, 31% used standard deviation of expression as the only feature, and 12% used mean and standard deviation as the only two features. Other expression features appeared in 21% of the models. These expression features tended to be correlated with mean expression across tissues (median r^2^=0.49). Disease association features appeared in 0.4% of the models.

**Table 4.**
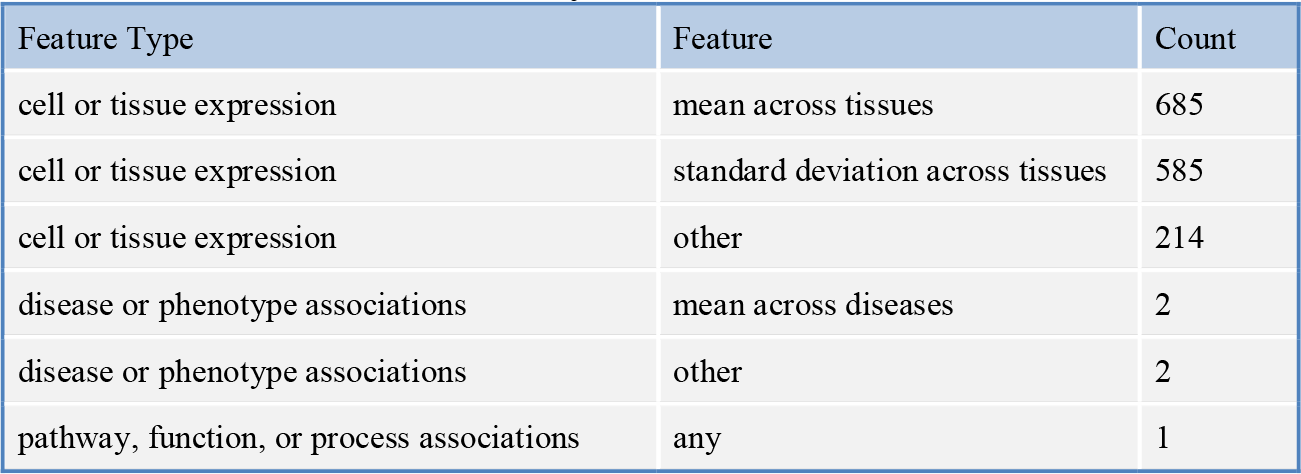
Number of train-test cycles in which feature was selected for the classifier.

#### Classifier performance

The classifier consistently had better than random performance in cross-validation (Fig 3, Table 5, Supplementary Table S5). The 2.5^th^, 50^th^, and 97.5^th^ percentiles for AUROC were 0.51, 0.57, and 0.61. For comparison, a random ordering of targets would yield an AUROC of 0.50. The receiver operating characteristic curve showed that there was no single cut-off that would provide satisfactory discrimination between successes and failures (Fig 3A). For an alternative view, we used kernel density estimation (59) to fit distributions of the probability of success predicted by the classifier for the successful, failed, and unlabeled targets (Fig 3B, Supplementary Table S1). The distributions for successes and failures largely overlapped, except in the tails.

**Fig 3.**
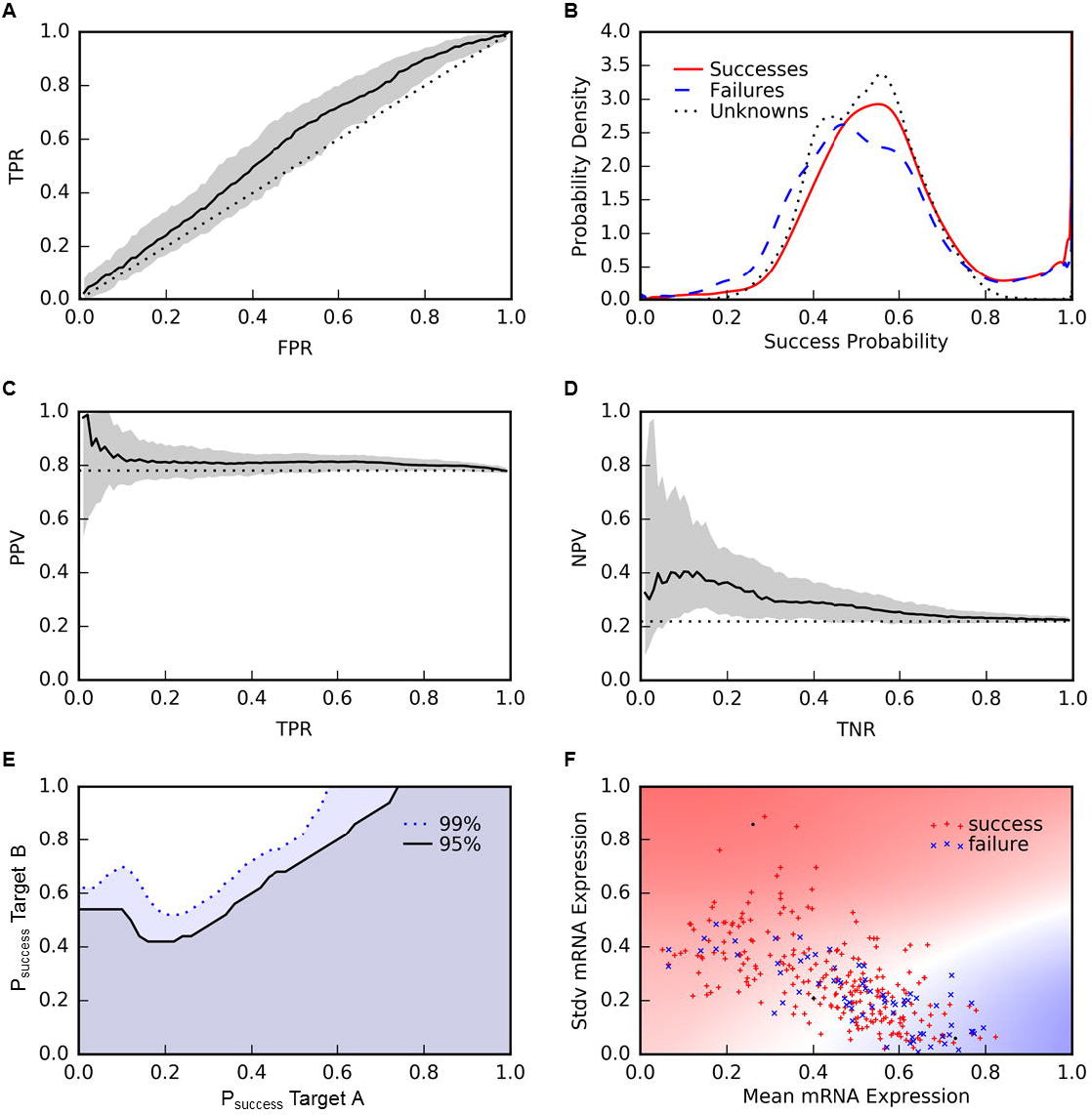
Classifier Performance. **(A)** Receiver operating characteristic (ROC) curve. The solid black line indicates the median performance across 200 repetitions of 5-fold cross-validation and the gray area indicates the range of the 2.5 and 97.5 percentiles. The dotted black line indicates the performance of random rankings. **(B)** Distributions of the probability of success predicted by the classifier for the successful, failed, and unlabeled targets. **(C)** Precision-recall curve for success predictions. **(D)** Precision-recall curve for failure predictions. **(E)** Pairwise target comparisons. For each pair of targets, we computed the fraction of repetitions of cross-validation in which Target B had a higher predicted probability of success greater than Target A. The heatmap illustrates this fraction, thresholded at 0.95 or 0.99, plotted as a function of the median predicted probabilities of success of two targets. The upper left region is where the classifier is 95% (above solid black line) or 99% (above dotted blue line) consistent in predicting greater probability of success of Target B than Target A. **(F)** Relationship between features and phase III outcomes. Heat map showing the projection of the predicted success probabilities onto the two dominant features selected for the classifier: mean expression across tissues and standard deviation of expression across tissues. Red, white, and blue background colors correspond to 1, 0.5, and 0 success probabilities. Red plusses and blue crosses mark the locations of the success and failure examples. It appears the model has learned that failures tend to have high mean expression and low standard deviation of expression across tissues, while successes tend to have low mean expression and high standard deviation of expression. The success and failure examples are not well separated, indicating that we did not discover enough features to fully explain why targets succeed or fail in phase III clinical trials.

**Table 5.**
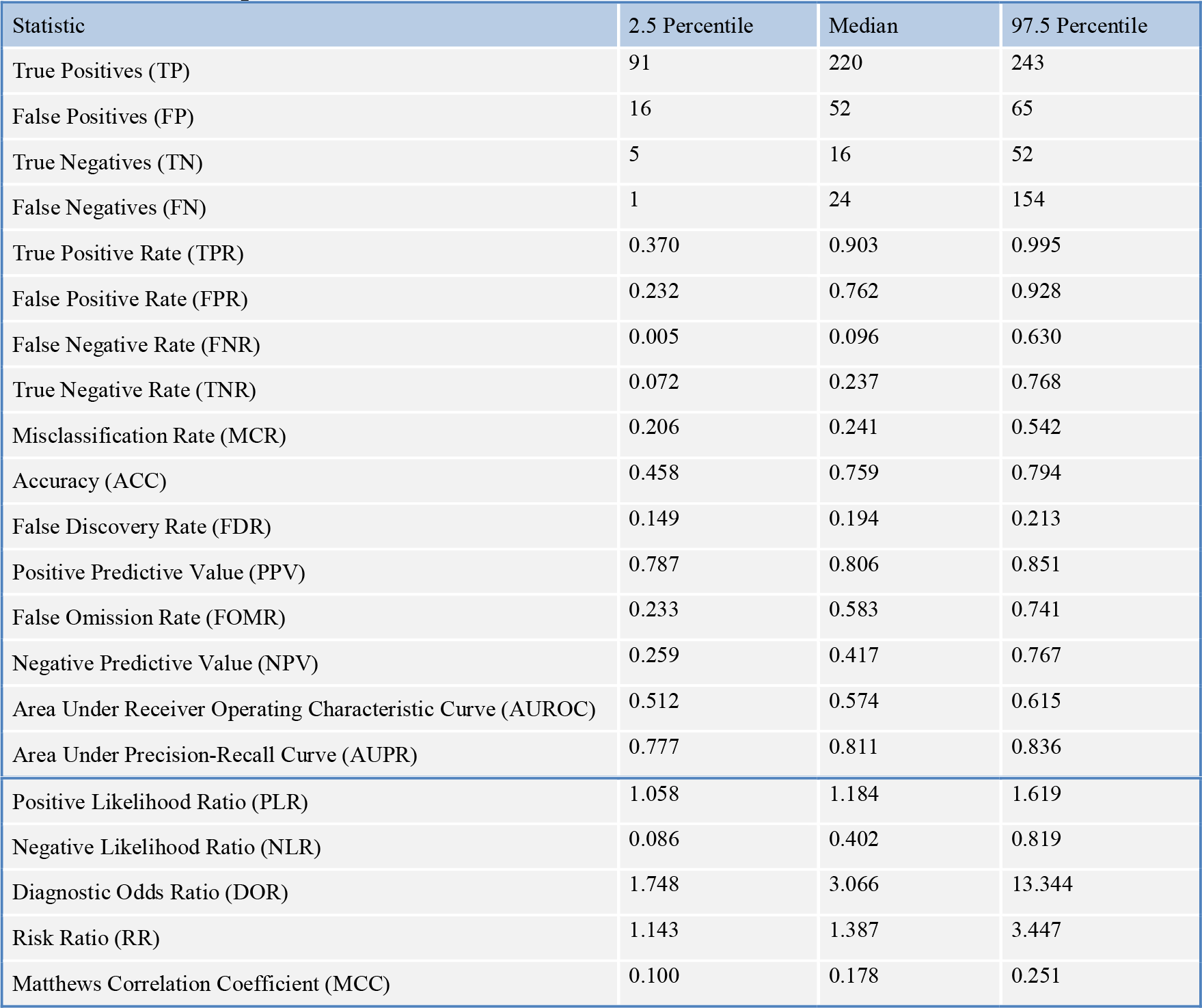
Classifier performance statistics.

We attempted to identify subsets of targets with high positive predictive value (PPV) or high negative predictive value (NPV). The median PPV rose as high as 0.99, but uncertainty in the PPV was so large that we could not be confident in identifying any subset of targets with a predicted success rate better than the historical 0.77 (Fig 3C). The median NPV rose to 0.40, roughly twice the historical failure rate of 0.23. Furthermore, at 0.40 median NPV, 99% of the cross-validation repetitions had an NPV greater than the historical failure rate (Fig 3D). Using this cut-off, we identified 943 unlabeled targets expected to be twice as likely to fail in phase III clinical trials as past phase III targets.

We reasoned that a more practical use of the classifier would be to make pair-wise comparisons among a short list of targets already under consideration for a therapeutic program. To assess the utility of the classifier for this purpose, for every pair of targets T_A_ and T_B_, we computed the fraction of cross-validation runs in which the classifier predicted greater probability of success for T_B_ than T_A_. We identified 67,270,678 target pairs (39%) with at least a 0.1 difference in median success probability where the classifier was 95% consistent in predicting greater probability of success for T_B_ than T_A_. The classifier was 99% consistent for 41528043 target pairs (24%). Requiring at least a 2-fold difference in median success probability between T_B_ and T_A_ reduced these counts to 2,730,437 target pairs (1.6%) at 95% consistency and 2,700,856 target pairs (1.6%) at 99% consistency. We visualized these results by plotting the 95% and 99% consistency fraction thresholds smoothly interpolated as a function of the median predicted probabilities of success of T_A_ and T_B_ (Fig 3E). For a median probability of success of T_A_ around 0.2, T_B_ must have a median probability of success of 0.5 or greater at the 99% threshold. For lower T_A_ success probabilities, the T_B_ success probability must be even higher because there is greater uncertainty about the low T_A_ probabilities. For higher T_A_ success probabilities, the T_B_ success probability at the 99% threshold increases steadily until a T_A_ success probability of about 0.6, where the T_B_ success probability reaches 1. For T_A_ success probabilities above 0.6, no targets are predicted to have greater probability of success with 99% consistency.

#### Feature interpretation

To interpret the relationship inferred by the classifier between target features and outcomes, we created a heatmap of the probability of success predicted by the classifier projected onto the two features predominantly selected for the model: mean expression and standard deviation of expression across tissues (Fig 3F). The probability of success was high in the subspace with low mean expression and high standard deviation of expression, and transitioned to low probability in the subspace with high mean expression and low standard deviation of expression. This trend appeared to be consistent with the distribution of the success and failure examples in the space.

## DISCUSSION

### Gene expression predicts phase III outcome

We searched over 150,000 target features from 67 datasets covering 16 feature types for predictors of target success or failure in phase III clinical trials (Table 1, Fig 1). We found several features significantly correlated with phase III outcome, robust to re-sampling, and generalizable across target classes (Table 2). To assess the usefulness of such features, we implemented a nested cross-validation routine to select features, train a classifier to predict the probability a target will succeed in phase III clinical trials, and estimate the stability and generalization performance of the model (Figs 2 and 3, Tables 3, 4, and 5). Ultimately, we found two features useful for predicting success or failure of targets in phase III clinical trials. Successful targets tended to have low mean mRNA expression across tissues and high standard deviation of mRNA expression across tissues (Fig 3F). These features were significant in multiple gene expression datasets, which increased our confidence that their relationship to phase III outcome was real, at least for the targets in our sample, which included only targets of selective drugs indicated for non-cancer diseases.

One interpretation of why the gene expression features were predictive of phase III outcome is that they are informative of the specificity of a target’s expression across tissues. A target with tissue specific expression would have a high standard deviation relative to its mean expression level. Tissue specific expression has been proposed by us and others as a favorable target characteristic in the past (4, 14, 60-62), but the hypothesis had not been evaluated empirically using examples of targets that have succeeded or failed in clinical trials. For a given disease, if a target is expressed primarily in the disease tissue, it is considered more likely that a drug will be able to exert a therapeutic effect on the disease tissue while avoiding adverse effects on other tissues. Additionally, specific expression of a target in the tissue affected by a disease could be an indicator that dysfunction of the target truly causes the disease.

The distribution of the success and failure examples in feature space (Fig 3F) partially supports the hypothesis that tissue specific expression is a favorable target feature. Successes were enriched among targets with low mean expression and high standard deviation of expression (tissue specific expression), and failures were enriched among targets with high mean expression and low standard deviation of expression (constitutive expression). However, it does not hold in general that, at any given mean expression level, targets with high standard deviation of expression tend to be more successful than targets with low standard deviation of expression. Nevertheless, our results encourage further investigation of the relationship between tissue specific expression and clinical trial outcomes. Deeper insight may be gleaned from analysis of gene expression features explicitly designed to quantify specificity of a target’s expression in the tissue(s) affected by the disease treated in each clinical trial.

### Caveats and limitations

Latent factors (variables unaccounted for in this analysis) could confound relationships between target features and phase III outcomes. For example, diseases pursued vary from target to target, and a target’s expression across tissues may be irrelevant for diseases where drugs can be delivered locally or for Mendelian loss-of-function diseases where treatment requires systemic replacement of a missing or defective protein. Also, clinical trial failure rates vary across disease classes (2). Although we excluded targets of cancer therapeutics from our analysis, we otherwise did not control for disease class as a confounding explanatory factor. Modalities (e.g. small molecule, antibody, antisense oligonucleotide, gene therapy, or protein replacement) and directions (e.g. activation or inhibition) of target modulation also vary from target to target and could be confounding explanatory factors or alter the dependency between target features and outcomes.

The potential issues described above are symptoms of the fact that our analysis (and any analysis of clinical trial outcomes) attempts to draw conclusions from a small (roughly 300 targets) and biased sample (63, 64). Latent factors such as target classes, disease classes, modalities, and directions of target modulation are not uniformly represented in the sample, yet correlations between target features and clinical trial outcomes likely depend on these factors. Unfortunately, attempts to stratify, match, or otherwise control for these factors are limited by the sample size. (The number of combinations of target class, disease class, modality, and direction of modulation exceeds the sample size.) We employed several tests to build confidence that our findings generalize across target classes, but did not address other latent factors. Consequently, we cannot be sure that conclusions drawn from this study apply equally to targets modulated in any direction, by any means, to treat any disease. For specific cases, expert knowledge and common sense should be relied upon to determine whether conclusions from this study (or similar studies) are relevant.

Another limitation is selection bias (63, 64). Targets of drugs are not randomly selected from the genome and cannot be considered representative of the population of all possible targets. Likewise, diseases treated by drugs are not randomly chosen; therefore, phase III clinical trial outcomes for each target cannot be considered representative of the population of all possible outcomes. Although we implemented tests to build confidence that our findings can generalize to new targets and new target classes, ultimately, no matter how we dissect the sample, a degree of uncertainty will always remain about the relevance of any findings for new targets that lack a representative counterpart in the sample.

Additionally, data processing and modeling decisions have introduced bias into the analysis. For example, we scored each target as successful or failed by its best outcome in all applicable (selective drug, non-cancer indication) phase III clinical trials. This approach ignores nuances. A target that succeeded in one trial and failed in all others is treated as equally successful as a target that succeeded in all trials. Also, the outcome of a target tested in a single trial is treated as equally certain as the outcome of a target tested in multiple trials. Representing target outcomes as success rates or probabilities may provide better signal for discovering features predictive of outcomes.

Another decision was to use datasets of features as we found them, rather than trying to reason about useful features that could be derived from the original data. Because of the breadth of data we interrogated, the effort and expertise necessary to hand engineer features equally well across all datasets exceeded our resources. Others have had success hand engineering features for similar applications in the past, particularly with respect to computing topological properties of targets in protein-protein interaction networks (18, 20, 21). This analysis could benefit from such efforts, potentially changing a dataset or feature type from yielding no target features correlated with phase III outcomes to yielding one or several useful features (22). On a related point, because we placed a priority on discovering interpretable features, we performed dimensionality reduction by averaging groups of highly correlated features and concatenating their (usually semantically related) labels. Dimensionality reduction by principal components analysis (65) or by training a deep auto-encoder (66) could yield more useful features, albeit at the expense of interpretability.

We cannot stress enough the importance of taking care not to draw broad conclusions from our study, particularly with respect to the apparent dearth of features predictive of target success or failure. We examined only a specific slice of clinical trial outcomes (phase III trials of selective drugs indicated for non-cancer diseases) summarized in a particular way (net outcome per target, as opposed to outcome per target-indication pair). Failure of a feature to be significant in our analysis should not be taken to mean it has no bearing on target selection. For example, prior studies have quantitatively shown that genetic evidence of disease association(s) is a favorable target characteristic (3, 36), but we did not find a significant correlation between genetic evidence and target success in phase III clinical trials. Our finding is consistent with the work of Nelson et al. (36), who investigated the correlation between genetic evidence and drug development outcomes at all phases and found a significant correlation overall and at all phases of development except phase III. As a way of checking our work, we applied our methods to test for features that differ between targets of approved drugs and the remainder of the druggable genome (instead of targets of phase III failures), and we recovered the finding of Nelson et al. that targets of approved drugs have significantly more genetic evidence than the remainder of the druggable genome (Supplementary Table S6). This example serves as a reminder to be cognizant of the domain of applicability of research findings. Though we believe we have performed a rigorous and useful analysis, we have shed light on only a small piece of a large and complex puzzle.

Advances in machine learning enable and embolden us to create potentially powerful predictive models for target selection. However, as described in the limitations, scarce training data are available, the data are far from ideal, and we must be cautious about building models with biased data and interpreting their predictions. For example, many features that appeared to be significantly correlated with phase III clinical trial outcomes in our primary analysis did not hold up when we accounted for target class selection bias. This study highlights the need for both domain knowledge and modeling expertise to tackle such challenging problems.

### Conclusion

Our analysis revealed several features that significantly separated targets of approved drugs from targets of drug candidates that failed in phase III clinical trials. This suggested that it is feasible to construct a model integrating multiple interpretable target features derived from Omics datasets to inform target selection. Only features derived from tissue expression datasets were promising predictors of success versus failure in phase III, specifically, mean mRNA expression and standard deviation of expression across tissues. Although these features were significant at a false discovery rate cut-off of 0.05, their effect sizes were too small to be useful for classification of the majority of untested targets, however, even a two-fold improvement in target quality can dramatically increase R&D productivity (67). We identified 943 targets predicted to be twice as likely to fail in phase III clinical trials as past phase III targets, and, therefore, should be flagged as having unfavorable expression characteristics. We also identified 2,700,856 target pairs predicted with 99% consistency to have a 2-fold difference in success probability, which could be useful for prioritizing short lists of targets with attractive disease relevance.

It should be noted that our analysis was not designed or powered to show that specific datasets or data types have no bearing on target selection. There are many reasons why a dataset may not have yielded any significant features in our analysis. In particular, data processing and filtering choices could determine whether or not a dataset or data type has predictive value. Also, latent factors, such as target classes, disease classes, modalities, and directions of target modulation, could confound or alter the dependency between target features and clinical trial outcomes. Finally, although we implemented tests to ensure robustness and generalizability of the target features significantly correlated with phase III outcomes, selection bias in the sample of targets available for analysis is a non-negligible limitation of this study and others of its kind. Nevertheless, we are encouraged by our results and anticipate deeper insights and better models in the future, as researchers improve methods for handling sample biases and learn more informative features.

## METHODS

### Data

#### Clinical Outcomes

We extracted data from Citeline’s Pharmaprojects database (38) (downloaded May 27, 2016), reformatting available XML data into a single tab-delimited form having one row for each asset (i.e. drug or drug candidate)/company combination. For each asset, known targets, identified with EntrezGene (68) IDs and symbols, and indications are reported. We obtained 107,120 asset-indication pairs and 37,211 asset-target pairs, correcting a single outdated EntrezGene ID, for SCN2A, which we updated from 6325 to 6326.

An overall pipeline status of each asset (e.g. “Launched”, “Discontinued”, “No Development Reported”) is reported in a single field (“Status”), and detailed information for each indication being pursued is dispersed throughout several other fields (e.g., “Key Event Detail”, “Overview”, etc.). While many assets have been tried against a single indication, and thus the status of the asset-indication pair is certain, the majority (N=61,107) of asset-indication pairs are for assets with multiple indications. For those pairs, we used a combination of string searching of these fields and manual review of the results to determine the likely pipeline location and status of each indication. For example, we excluded efforts where a trial of an asset was reported as planned, but no further information was available. Asset-indication pairs were thus assigned a status of Successful (“Launched”, “Registered”, or “Pre-registration”), Failed (“Discontinued”, “No Development Reported”, “Withdrawn”, or “Suspended”), or In Progress, consisting of 9,337, 72,269 and 25,159 pairs, respectively. We then used the pipeline location to assign each asset-indication pair to one of 10 outcomes: Succeeded, In Progress-Preclinical, In Progress-Phase I, In Progress-Phase II, In Progress-Phase III, Failed-Preclinical, Failed-Phase I, Failed-Phase II, Failed-Phase III, and Failed-Withdrawn. We discarded indications which were diagnostic in nature or unspecified, mapping the remainder to Medical Subject Headings (MeSH) (69). We also observed that only 24% of the failures reported in Pharmaprojects are clinical failures, suggesting a clinical success rate of nearly 35%, much higher than typically cited (67).

We joined the list of asset-indication-outcome triples with the list of asset-target pairs to produce a list of asset-target-indication-outcome quadruples. We then filtered the list to remove: 1) assets with more than one target, 2) non-human targets, 3) cancer indications (indications mapped to MeSH tree C04), and 4) outcomes labeled as In Progress at any stage or Failed prior to Phase III. We scored the remaining targets (N=331) as Succeeded (N=259), if the target had at least one successful asset remaining in the list, or Failed (N=72), otherwise.

#### Target Features

We obtained target features from the Harmonizome (39), a recently published collection of features of genes and proteins extracted from over 100 Omics datasets. We downloaded (on June 30, 2016) a subset of Harmonizome datasets that were in the public domain or GSK had independently licensed (Table 1). Each dataset was structured as a matrix with genes labeling the rows and features such as diseases, phenotypes, tissues, and pathways labeling the columns. Genes were identified with EntrezGene IDs and symbols, enabling facile integration with the clinical outcome data from Pharmaprojects. Some datasets were available on the Harmonizome as a “cleaned” version and a “standardized” version. In all instances, we used the cleaned version, which preserved the original data values (e.g. gene expression values), as opposed to the standardized version, in which the original data values were transformed into scores indicating relative strengths of gene-feature associations intended to be comparable across datasets. The data matrices were quantitative and filled-in (e.g. gene expression measured by microarray), quantitative and sparse (e.g. protein expression measured by immunohistochemistry), or categorical (i.e. binary) and sparse (e.g. pathway associations curated by experts). We standardized quantitative, filled-in features by subtracting the mean and then dividing by the standard deviation. We scaled quantitative, sparse features by dividing by the mean. We included the mean and standard deviation calculated along the rows of each dataset as additional target features. We excluded features that had fewer than three non-zero values for the targets with phase III clinical trial outcomes. The remaining features, upon which our study was based, have been deposited at https://github.com/arouillard/omic-features-successful-targets.

### Dimensionality Reduction

Our goals in performing dimensionality reduction were to identify groups of highly correlated features, avoid excessive multiple hypothesis testing, and maintain interpretability of features. For each dataset, we computed pair-wise feature correlations (r) using the Spearman correlation coefficient (70-72) for quantitative, filled-in datasets, and the cosine coefficient (71, 72) for sparse or categorical datasets. We thresholded the correlation matrix at r^2^=0.5 (for the Spearman correlation coefficient, this corresponds to one feature explaining 50% of the variance of another feature, and for the cosine coefficient, this corresponds to one feature being aligned within 45 degrees of another feature) and ordered the features by decreasing number of correlated features. We created a group for the first feature and its correlated features. If the dataset mean was included in the group, we replaced the group of features with the dataset mean. Otherwise, we replaced the group of features with the group mean and assigned it the label of the first feature (to indicate that the feature represents the average of features correlated with the first feature), while also retaining a list of the labels of all features included in the group. We continued through the list of features, repeating the grouping process as described for the first feature, except excluding features already assigned to a group from being assigned to a second group.

### Feature Selection

We performed permutation tests (40, 41) to find features with a significant difference between successful and failed targets. We used permutation testing in order to apply the same significance testing method to all features. The features in our collection had heterogeneous shapes of their distributions and varying degrees of sparsity, and therefore no single parametric test would be appropriate for all features. Furthermore, individual features frequently violated assumptions required for parametric tests, such as normality for the t-test (for continuous-valued features) or having at least five observations in each entry of the contingency table for the Chi-squared test (for categorical features). For each feature, we performed 10^5^ success/failure label permutations to obtain a null distribution for the difference between the means of successful and failed targets, and then calculated an empirical two-tailed p-value as the fraction of permutations that yielded a difference between means at least as extreme as the actual observed difference. We used the Benjamini-Yekutieli method (42) to correct for multiple hypothesis testing within each dataset and accepted features with corrected p-values less than 0.05 as significantly correlated with phase III clinical trial outcomes, thus controlling the false discovery rate at 0.05 within each dataset.

### Feature Robustness and Generalizability

#### Robustness to sample variation

We used bootstrapping (53, 54) to investigate how robust our significance findings were to variation in the success and failure examples. We created a bootstrap sample by sampling with replacement from the original set of examples to construct an equal sized set of examples. For each dataset that yielded significant features in our primary analysis, we repeated the analysis on the bootstrap sample and recorded whether the features were still significant at the aforementioned 0.05 false discovery rate cut-off. We performed this procedure on 1000 bootstrap samples and quantified the replication probability (55) of each feature as the fraction of bootstraps showing a significant correlation between the feature and phase III clinical trial outcomes. We accepted features with replication probabilities greater than 0.8 (55) as robust to sample variation.

#### Robustness to target class variation

We tested if any of the significance findings depended upon the presence of targets from a single target class in our sample. We obtained target class labels (i.e. gene family labels) from the HUGO Gene Nomenclature Committee (56) (downloaded April 19, 2016) and created binary features indicating target class membership. Using the same permutation testing and multiple hypothesis testing correction methods described above for feature selection, we tested if any target classes were significantly correlated with phase III clinical trial outcomes. Then, we tested if the significant target classes were correlated with any significant features. Such features might be correlated with clinical outcome only because they are surrogate indicators for particular target classes that have been historically very successful or unsuccessful, as opposed to the features being predictors of clinical outcome irrespective of target class. To test this possibility, we performed a bootstrapping procedure as described above, except did not allow examples from target classes correlated with clinical outcome to be drawn when re-sampling. Thus, the modified bootstrapping procedure provided replication probabilities conditioned upon missing information about target classes correlated with clinical outcome. We accepted features with replication probabilities greater than 0.8 as robust to target class variation.

#### Generalization across target classes

We implemented a modified permutation test, inspired by the approach of Epstein et al. (57) to correct for confounders in permutation testing, to select features correlated with phase III clinical trial outcomes while controlling for target class as a confounding explanatory factor. In the modified permutation test, success/failure labels were shuffled only within target classes, so the sets of null examples had the same ratios of successes to failures within target classes as in the set of observed examples. Consequently, features had to correlate with clinical outcome within multiple classes to be significant, while features that discriminated between classes would not be significant. We performed bootstrapping as described previously to obtain replication probabilities for the significant features, in this case conditioned upon including target class as an explanatory factor. We accepted features with replication probabilities greater than 0.8 as generalizable across target classes represented in the sample.

### Clinical Outcome Classifier

We trained a classifier to predict target success or failure in phase III clinical trials, using a procedure like the above for initial feature selection, then using cross-validation to select a model type (Random Forest or logistic regression) and subset of features useful for prediction. We used an outer cross-validation loop with 5-folds repeated 200 times, yielding a total of 1000 train-test cycles, to estimate the generalization performance and stability of the feature selection and model selection procedure (58). Each train-test cycle had five steps: 1) splitting examples into training and testing sets, 2) univariate feature selection on the training data, 3) aggregation of significant features from different datasets into a single feature matrix, 4) model selection and model-based (multivariate) feature selection on the training data, and 5) evaluation of the classifier on the test data.

#### Step 2: Univariate feature selection

Beginning with the non-redundant features obtained from dimensionality reduction, we performed modified permutation tests to find features with a significant difference between successful and failed targets in the training examples. As described above, for the modified permutation test, success/failure labels were shuffled only within target classes. This was done to control for target class as a confounding factor that might explain correlations between phase III outcomes and features. For each feature, we performed 10^4^ success/failure label permutations and calculated an empirical two-tailed p-value. We corrected for multiple hypothesis testing within each dataset and accepted features with corrected p-values less than 0.05.

#### Step 3. Feature aggregation

Significant features from different datasets, each having different target coverage, had to be aggregated into a single feature matrix prior to training a classifier. When features from many datasets were aggregated, we found that the set of targets with no missing data across all features could become very small. To mitigate this, we excluded features from non-human datasets and small datasets (fewer than 2,000 genes). We also excluded features from the Allen Brain Atlas human brain expression atlas, unless there were no other significant features, because we noticed it had poor coverage of targets with phase III outcomes (287) compared to other expression atlases, such as BioGPS (320), GTEx (328), and HPA (314), which almost always yielded alternative significant expression-based features. After aggregating features into a single matrix, we min-max scaled the features so that features from different datasets would have the same range of values (from 0 to 1).

To reduce redundancy in the aggregated feature matrix, we grouped features as described for the primary analysis. We used the cosine coefficient to compute pair-wise feature correlations because some features were sparse. Instead of replacing groups of correlated features with the group mean, we selected the feature in each group that was best correlated with phase III outcomes, because we preferred not to create features derived from multiple datasets.

#### Step 4. Model selection and model-based feature selection

We hypothesized that a Random Forest classifier (73) would be a reasonable model choice because the Random Forest model does not make any assumptions about the distributions of the features and can seamlessly handle a mixture of quantitative, categorical, filled-in, or sparse features. Furthermore, we expected each train-test cycle to yield only a handful of significant features. Consequently, we would have 10- to 100-fold more training examples than features and could potentially afford to explore non-linear feature combinations. We also trained logistic regression classifiers and used an inner cross-validation loop (described below) to choose between Random Forest and logistic regression for each train-test cycle of the outer cross-validation loop. We used the implementations of the Random Forest and logistic regression classifiers available in the Scikit-learn machine learning package for Python. To correct for unequal class sizes during training, the loss functions of these models weighted the training examples inversely proportional to the size of each example’s class.

We performed incremental feature elimination with an inner cross-validation loop to 1) choose the type of classifier (Random Forest or logistic regression) and 2) choose the smallest subset of features needed to maximize the performance of the classifier. First, we trained Random Forest and logistic regression models using the significant features aggregated in Step 2, performing 5-fold cross-validation repeated 20 times to obtain averages for the area under the receiver operating characteristic curve (AUROC) and area under the precision recall curve (AUPR). We also obtained average feature importance scores from the Random Forest model. Next, we eliminated the feature with lowest importance score and trained the models using the reduced feature set, performing another round of 5-fold cross-validation repeated 20 times to obtain AUROC, AUPR, and feature importance scores. We continued eliminating features then obtaining cross-validation performance statistics and feature importance scores until no features remained. Then, we found all models with performance within 95% of the maximum AUROC and AUPR. If any logistic regression models satisfied this criterion, we selected the qualifying logistic regression model with fewest features. Otherwise, we selected the qualifying Random Forest model with fewest features.

#### Step 5. Classifier evaluation

For each train-test cycle, after selecting a set of features and type of model (Random Forest or logistic regression) in Step 4, we re-fit the selected model to the training data and predicted success probabilities for targets in the test set as well as unlabeled targets. For each round of 5-fold cross-validation, we computed the classifier’s receiver operating characteristic curve, precision-recall curve, and performance summary statistics, including the true positive rate, false positive rate, positive predictive value, negative predictive value, and Matthews correlation coefficient.

We computed distributions of the log odds ratios predicted by the classifier (log of the ratio of the predicted probability of success over the probability of failure) for the successful, failed, and untested (unlabeled) targets, aggregating predicted probabilities from the 200 repetitions of 5-fold cross-validation. Histograms of the log odds ratios for the three groups of targets were roughly bell-shaped, so we fit the distributions using kernel density estimation (59) with a Gaussian kernel and applied Silverman’s rule for the bandwidth. We transformed the fitted distributions from a function of log odds ratio to a function of probability of success using the rule pdf(x) = pdf(y)*|dy/dx|.

We created a heatmap of the probability of success predicted by the classifier projected onto the two dominant features in the model: mean mRNA expression across human tissues and standard deviation of mRNA expression across human tissues. We examined the heatmap to interpret the classifier’s decision function and assess its plausibility.

To more concretely assess the usefulness of the classifier, we found the probability cut-off corresponding to the maximum median positive predictive value and determined the number of unlabeled targets predicted to succeed at that cut-off. Likewise, we found the probability cut-off corresponding to the maximum median negative predictive value and determined the number of unlabeled targets predicted to fail at that cut-off. We also created a heatmap illustrating the separation needed between the median predicted success probabilities of two targets in order to be confident that one target is more likely to succeed than the other. This heatmap was created by calculating the fraction of times Target B had greater probability of success than Target A across the 200 repetitions of 5-fold cross-validation, for all pairs of targets.

### Implementation

Computational analyses were written in Python 3.4.5 and have the following package dependencies: Fastcluster 1.1.20, Matplotlib 1.5.1, Numpy 1.11.3, Requests 2.13.0, Scikit-learn 0.18.1, Scipy 0.18.1, and Statsmodels 0.6.1. Code, documentation, and data have been deposited on GitHub at https://github.com/arouillard/omic-features-successful-targets.

## ACKNOWLEDGMENTS

Many thanks to Dr. Subhas Chakravorty for assisting with access to and processing of the Pharmaprojects data and to Dr. David Cooper for helpful direction regarding nested cross-validation.

## SUPPORTING INFORMATION

**S1. Supplementary Table S1.** List of targets with their phase III outcome labels and predicted success probabilities for 200 cross-validation repetitions.

**S2. Supplementary Table S2.** List of non-redundant features with their similar features and p-values from the basic permutation test.

**S3. Supplementary Table S3.** List of classifier attributes (selected features, selected model type, and test performance) for 1000 train-test cycles.

**S4. Supplementary Table S4.** Comparison of inner cross-validation loop AUROC and AUPR values between Random Forest and logistic regression models for 1000 train-test cycles.

**S5. Supplementary Table S5.** List of classifier test performance statistics for 200 cross-validation repetitions.

**S6. Supplementary Table S6.** Cases illustrating how the significance of genetic evidence (and likely other types of evidence) as a predictor of target success depends on which targets are compared.

